# Perirhinal cortex supports object perception by integrating over visuospatial sequences

**DOI:** 10.1101/2023.09.07.556737

**Authors:** Tyler Bonnen, Anthony D. Wagner, Daniel L.K. Yamins

## Abstract

Perception unfolds across multiple timescales. For humans and other primates, many object-centric visual attributes can be inferred ‘at a glance’ (i.e., given *<*200ms of visual information), an ability supported by ventral temporal cortex (VTC). Other perceptual inferences require more time; to determine a novel object’s identity, we might need to represent its unique configuration of visual features, requiring multiple ‘glances.’ Here we evaluate whether perirhinal cortex (PRC), downstream from VTC, supports object perception by integrating over such visuospatial sequences. We first compare human visual inferences directly to electrophysiological recordings from macaque VTC. While human performance ‘at a glance’ is approximated by a linear readout of VTC, participants radically outperform VTC given longer viewing times (i.e., *>*200ms). Next, we leverage a stimulus set that enables us to characterize PRC involvement in these temporally extended visual inferences. We find that human visual inferences ‘at a glance’ resemble the deficits observed in PRC-lesioned human participants. Not surprisingly, by measuring gaze behaviors during these temporally extended viewing periods, we find that participants sequentially sample task-relevant features via multiple saccades/fixations. These patterns of visuospatial attention are both reliable across participants and necessary for PRC-dependent visual inferences. These data reveal complementary neural systems that support visual object perception: VTC provides a rich set of visual features ‘at a glance’, while PRC is able to integrate over the sequential outputs of VTC to support object-level inferences.

## 1 Introduction

There is temporal structure in how we perceive the world. Many visual attributes can be inferred ‘at a glance’ (Potter, 1975). Given just 100ms of visual information, humans and other primates can determine behaviorally relevant stimulus properties, such as object shape and category membership (Greene and Oliva, 2009; Potter and Levy, 1969). Ventral temporal cortex (VTC) supports these rapid visual inferences through successive, hierarchical transformations over retinal inputs (DiCarlo et al., 2012; Grill-Spector and Weiner, 2014). However, not all visual inferences are possible ‘at a glance’ (Findlay and Gilchrist, 2003). This is due, in part, to biological constraints on the primate visual system (Van Essen and Anderson, 1990). High-acuity visual information is only maintained at the central visual field (i.e., the fovea) and, as a consequence, humans typically shift gaze location three times per second (Hirsch and Curcio, 1989; Liversedge and Findlay, 2000). It has been proposed that these ‘visual routines’ are an essential component of primate vision: feedforward processes automatically extract visual information, which is then flexibly recombined to support goal-relevant behaviors (Ullman, 1987). While a great deal is known about the feedforward component of primate vision, less is known about the neural substrates that enable us to integrate across visuospatial sequences.

Perirhinal cortex (PRC) may play a critical role in integrating across visual sequences. Decades of lesion studies in humans and monkeys have demonstrated that PRC enables visual inferences not supported by VTC alone (e.g., Barense et al., 2007; Bussey and Saksida, 2002; Lee et al., 2005; Murray and Bussey, 1999). Theoretical (Murray and Wise, 2012) and computational (Cowell, 2012) accounts of perirhinal function indicate that while VTC contains low-level visual representations such as texture and part-based-shape features, PRC is unique in it’s ability to represent the ‘configural’ properties of objects by flexibly combining these features (Saksida and Bussey, 2010). While these results are often considered in terms of the *representations* supported by PRC, recent data—including observations about temporal dynamics (Bonnen et al., 2021) and gaze patterns (Erez et al., 2013)—highlight the need to characterize the *algorithmic* basis of perirhinal function. Considering this cumulative body of evidence, it is possible that PRC enables supra-VTC performance by integrating over the sequential outputs from high-level visual cortex. That is, MTC might support visual inferences not possible ‘at a glance.’

PRC is situated at the apex of VTC, such that it receives inputs from high-level visual cortex (Suzuki and Amaral, 2004). At the circuit-level, PRC is a ‘mesocortical’ structure (Brodmann, 1909; Vogt and Vogt, 1919), which is a laminar intermediate between six-layer isocortex and three-layer allocortex, providing a structural transition between these computational motifs. Functionally, PRC has many unique properties that distinguish it from earlier stages of visual processing (Miyashita, 2019). Here we simply note that, in contrast to the relatively transient neural responses in VTC (Xiao et al., 2024), the temporal properties of PRC neurons (e.g., recurrent and persistent firing dynamics) enable PRC to integrate information across much longer timescales (Kholodar-Smith et al., 2008a; Kholodar-Smith et al., 2008b; Leung et al., 2006). Given that lesion data typically include damage to PRC and PRC-adjacent entorhinal cortex (ERC), the available data are not able to isolate PRC’s functional role from all other MTC structures. Nonetheless, PRC’s structural and functional properties of PRC suggest that this neural substrate is ideally suited to integrate over the sequential outputs from VTC.

Here we characterize the complementary roles that VTC and PRC play in visual object perception using a combination of lesion, electrophysiological, computational, and psychophysics data. By comparing human performance to electrophysiological responses from high-level visual cortex, we first demonstrate that rapid visual inferences reflect VTC supported performance. However, with increased viewing time participants radically outperform VTC. These temporal relationships are also evident when comparing human performance to ‘VTC-like’ computational models: our rapid visual inferences are approximated by task-optimized convolutional neural networks (CNNs), but we radically outperform these models with more time. This enables us to use these models as proxies for VTC in subsequent experiments. Next, we implicate PRC in temporally extended visual behaviors by evaluating PRC-lesioned and -intact performance. We first validate our previous findings in this novel dataset, observing that human accuracy corresponds with VTC model performance at restricted viewing times, but exceeds VTC model performance with longer viewing times. Critically, we find that time restricted/unrestricted performance mirrors the differences between PRC-lesioned/-intact participants: neurologically intact ‘time restricted’ participant behaviors resemble the performance of PRC-lesioned participants. Finally, we use a series of eye-tracking experiments to reveal the sequential nature of information processing during these temporally extended, PRC-dependent visual inferences.

## 2 Comparing human performance to neural responses from VTC

### 2.1 Experimental designs used to isolate temporal information processing

We first outline the logic behind our experimental approach, then elaborate on the specific design choices and tasks we have employed. We examine human performance in two temporal regimes: ‘time restricted’ (i.e., ‘at a glance’, given less than 200ms of visual information), and ‘time unrestricted’ (i.e., ‘self-paced’). We draw this partition at 200ms because this is roughly the amount of time needed to initiate an eye movement (i.e., a saccade) in response to a stimulus (Leigh and Zee, 2015). When a stimulus is presented for just 100ms, for example, participants are unable to sequentially sample the image; performance must rely on visually processing ‘at a glance.’ Conversely, when stimuli are presented for longer durations (e.g., 1000ms) participants are able to collect high-acuity visual information from multiple locations via eye movements. We leverage two experimental designs to examine these temporal dynamics. First, we use a 3-way concurrent visual discrimination task (schematized in Fig. 3) commonly used to evaluate the role of PRC in perception (e.g., Barense et al., 2007; Buckley et al., 2001; Bussey et al., 2002). This design enables us to determine visual inferences that are possible with unlimited viewing time: on each trial, participants are presented with three images and must identify the image that does not match the other two in terms of object identity (i.e., the ‘oddity’). To evaluate the effect of time restriction on performance, we use a match-to-sample visual discrimination paradigm (schematized in Fig. 3), common in experiments used to evaluate VTC function (DiCarlo and Cox, 2007; DiCarlo et al., 2012; Majaj et al., 2015). In each match-to-sample trial, participants first view a ‘sample’ image, which disappears after 100ms, then two images are presented on the subsequent screen. Participants are instructed to select the image that ‘matches’ the previous image on the sample screen. We administer these experiments to human participants online (*N* = 474). We note that are many differences between these two designs; we address these issues with control experiments later in this section.

### 2.2 Estimating VTC/model performance directly from experimental stimuli

To compare to human visual abilities to VTC-supported performance, we draw from previously collected stimuli and electrophysiological data (Majaj et al., 2015. This stimulus set consists of 5760 images with associated population-level electrophysiological responses recorded from primate V4 and IT. We use a subset of these images (Methods: Generating a novel stimulus set from Majaj et al., 2015) to construct odd-one-out tasks (examples images from several trials can be found in Fig. 2). To estimate VTC supported performance on the stimuli above, we use a modified leave-one-out cross validation strategy. For a given sample_*ij*_ trial we construct a random combination of three-way oddity tasks to be used as training data; we sample without replacement from the pool of all images of object_*i*_ and object_*j*_, excluding only those three stimuli that were present in sample_*ij*_. This yields ‘pseudo oddity experiments’ where each trial contains two typical objects and one oddity that have the same identity as the objects in sample_*ij*_ and are randomly configured (different viewpoints, different backgrounds, different orders).

**Figure 1:**
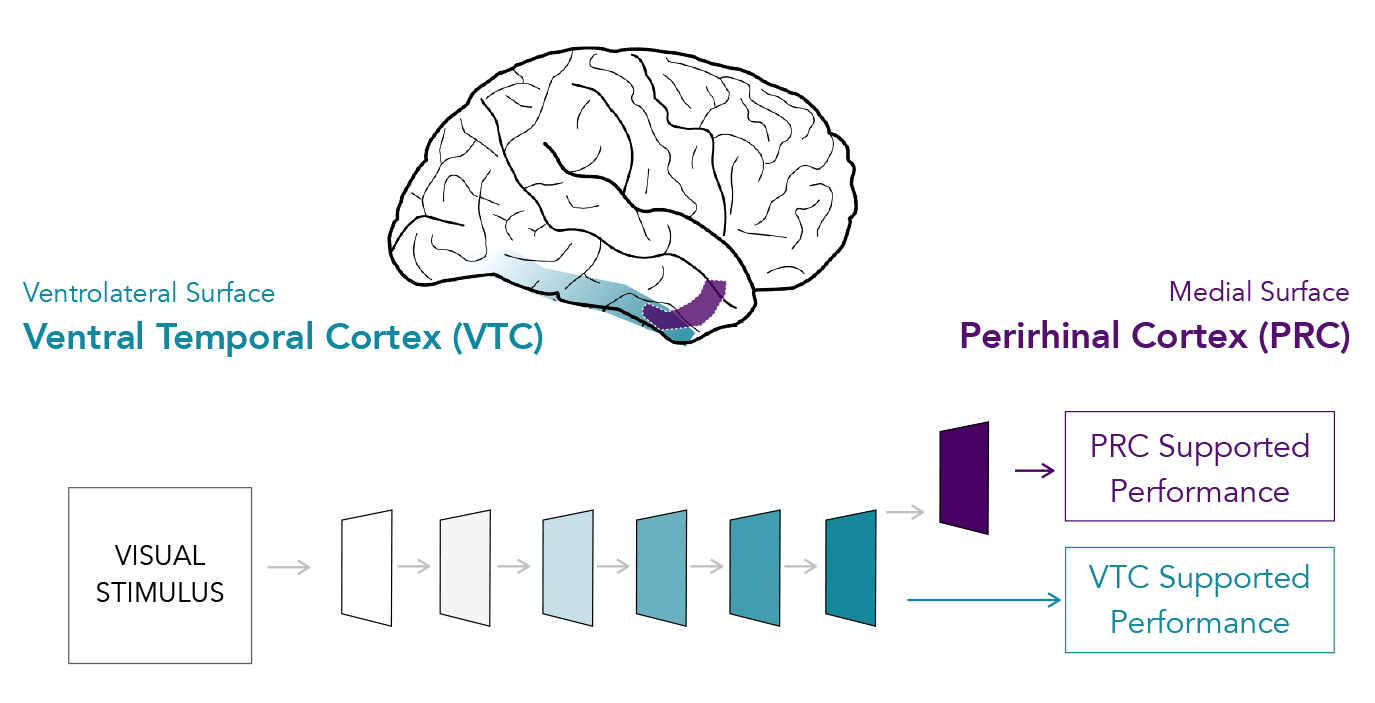
Anatomical relationship between ventral temporal cortex and perirhinal cortex. Ventral temporal cortex (VTC) is a hierarchical network of neural structures on the lateral and ventral surface of the temporal lobe (green). These structures have been shown to support rapid visual inferences through (largely) feedforward processing. Medial temporal cortex (MTC) is situated on the medial surface of the temporal lobe and is composed of numerous structures. Here, we visualize perirhinal cortex (PRC, purple), a MTC structure situated at the apex of VTC, such that it receives inputs from high-level VTC. While PRC has historically been studied in relation to memory-related behaviors, it is increasingly recognized for its role in other cognitive functions, including perception.

**Figure 2:**
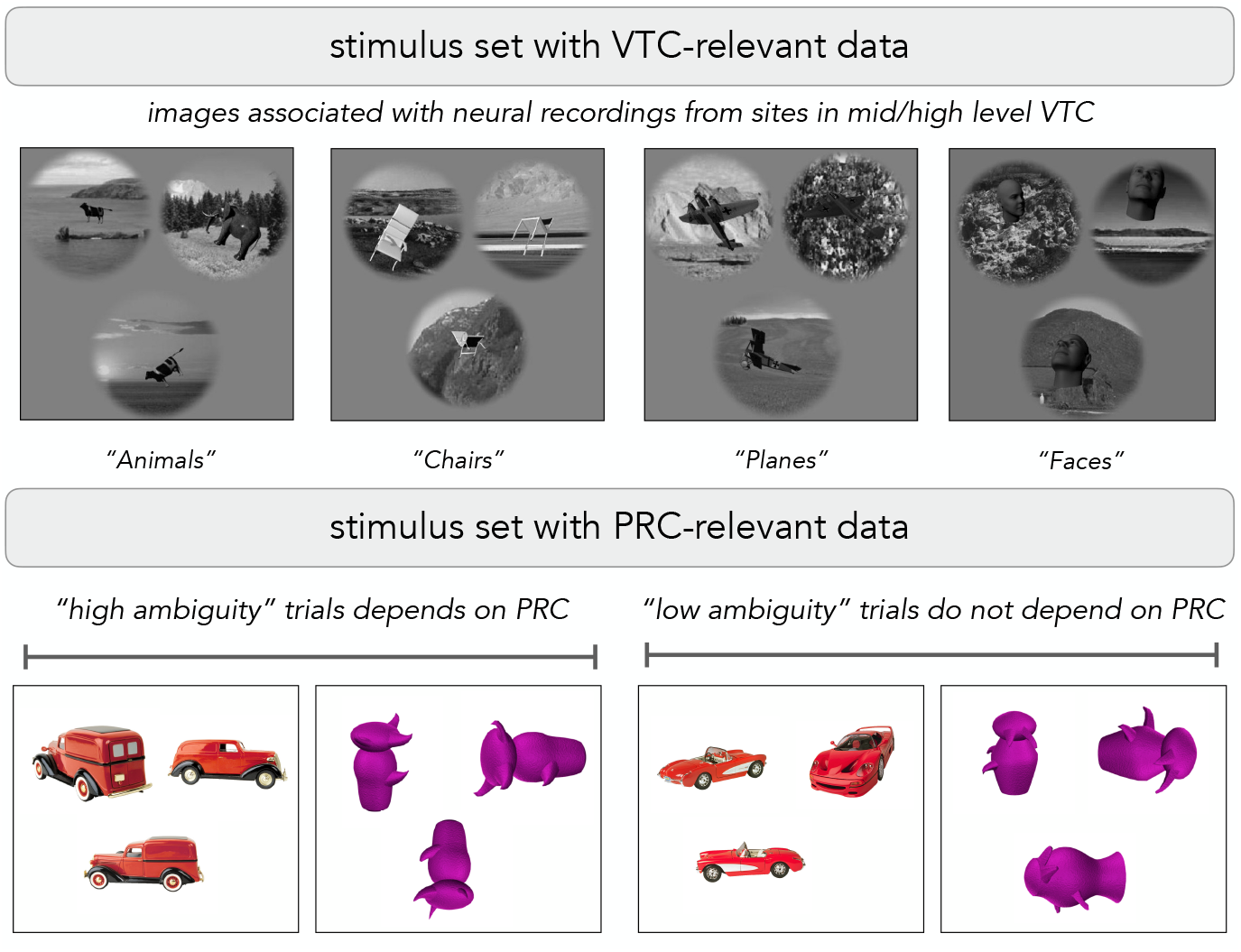
textbfExample stimuli from two datasets related to VTC and PRC function. We draw from stimuli and associated data collected by two previous experiments, enabling us to relate the psychophysics experiments in the current study to the behaviors supported by VTC and PRC. Majaj et al., 2015 presented a set of experimental stimuli (top) to two awake, head-fixed macaques, and collected neural recordings from mid (V4) and high (IT) level regions of ventral temporal cortex. We use these individual images to generate both ‘oddity’ (visualized here, top row) and sequential match to sample tasks. This enables us to compare human performance directly to neural recordings collected from high-level visual cortex. Barense et al., 2007 used a set of experimental stimuli (bottom) in a series of ‘oddity’ tasks administered to PRC-lesioned participants, as well as a group of age and IQ matched controls. On ‘high ambiguity’ trials (bottom left) PRC-lesioned participants were significantly impaired, while there was no such impairment on ‘low ambiguity’ trials (bottom right). We use these individual images to generate both ‘oddity’ (visualized here, bottom row) and sequential match to sample tasks. This enables us to evaluate human performance on trials that we know depend on PRC.

**Figure 3:**
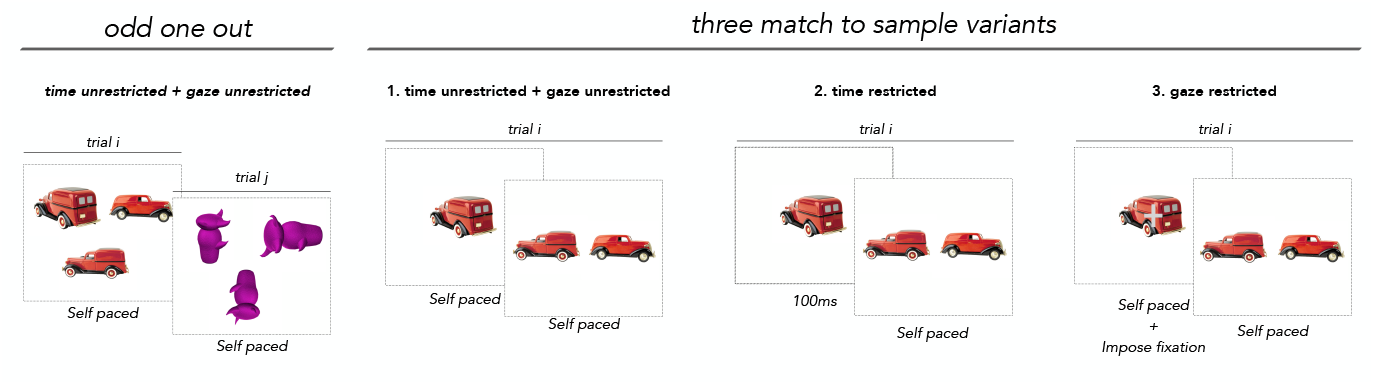
Visualizing the four experimental designs used in this manuscript. We use four complementary experimental designs in this study. Odd one out tasks (far left) enable us to estimate an upper bound of human visual inferences in the absence of working memory demands, while match to sample tasks (three on right) provide experimental control around temporal and foveal dynamics.

These ‘pseudo oddity experiments’ are used as training data. For each of these images, we identify neural responses from high-level visual cortex. We then train an L2 regularized linear classifier to identify the oddity across all (*N* = 52) trials in this permutation of pseudo oddity experiments generated for sample_*ij*_. After learning this weighted, linear readout, we evaluate the classifier on the neural responses to sample_*ij*_. This results in a prediction which is binarized into a single outcome *{* 0 | 1*}*, either correct or incorrect. We repeat this protocol across 100 random sample_*ij*_s, and average across them, resulting in a single estimate of VTC supported performance for each pair_*ij*_. Additionally, we estimate the performance from a computational proxy for VTC using an analogous protocol: instead of using neural responses from VTC, we use model responses from a layer that predicts VTC responses (Methods: Determining VTC model performance on Majaj et al., 2015).

### 2.3 VTC matches human performance ‘at a glance,’ but not at longer timescales

We compare time restricted and unrestricted human visual inferences directly to electrophysiological recordings collected from the macaque. This dataset (Majaj et al., 2015) contains neural responses from high-level visual cortex (i.e., inferior temporal cortex, within VTC in the macaque) to stimuli from semantic categories (e.g., Animals, Chairs, Planes, Faces; examples in Fig. 2). With a subset of these images (Methods: Generating a novel stimulus set from Majaj et al., 2015) we use a modified leave-one-out cross validation strategy, enabling us to estimate the item-level (*N* = 32) performance supported by a linear readout of high-level visual cortex (Methods: Estimating VTC supported performance on stimuli from Majaj et al., 2015). We administer these same stimuli to human participants (Methods: Online human data collection). We observe a striking correspondence between time restricted human performance (estimated using a sequential match to sample task, *N* = 107) and a linear readout of VTC (item-level analysis, *β* = 0.87, *F* (1, 30) = 8.34, *P* = 3×10^*−*9^; Fig. 4, left, green). There is not a significant difference between human accuracy and VTC supported performance (item-level paired ttest *β* = *−*0.02, *t*(31) = *−*0.85, *P* = 0.40). However, with unrestricted viewing time (estimated using the simultaneous visual discrimination task, *N* = 297), participants substantially outperform a linear readout of VTC (Fig. 4, left (purple): item-level paired ttest *β* = .24, *t*(31) = 9.50, *P* = 1×10^*−*10^). We observe a significant interaction between human and VTC supported performance as a function of stimulus presentation time (*β* = 0.54, *F* (3, 60) = 5.25, *P* = 2 × 10^*−*6^). These data indicate that human performance ‘at a glance’ reflects a linear readout of VTC, while humans substantially outperform VTC given longer viewing times.

**Figure 4:**
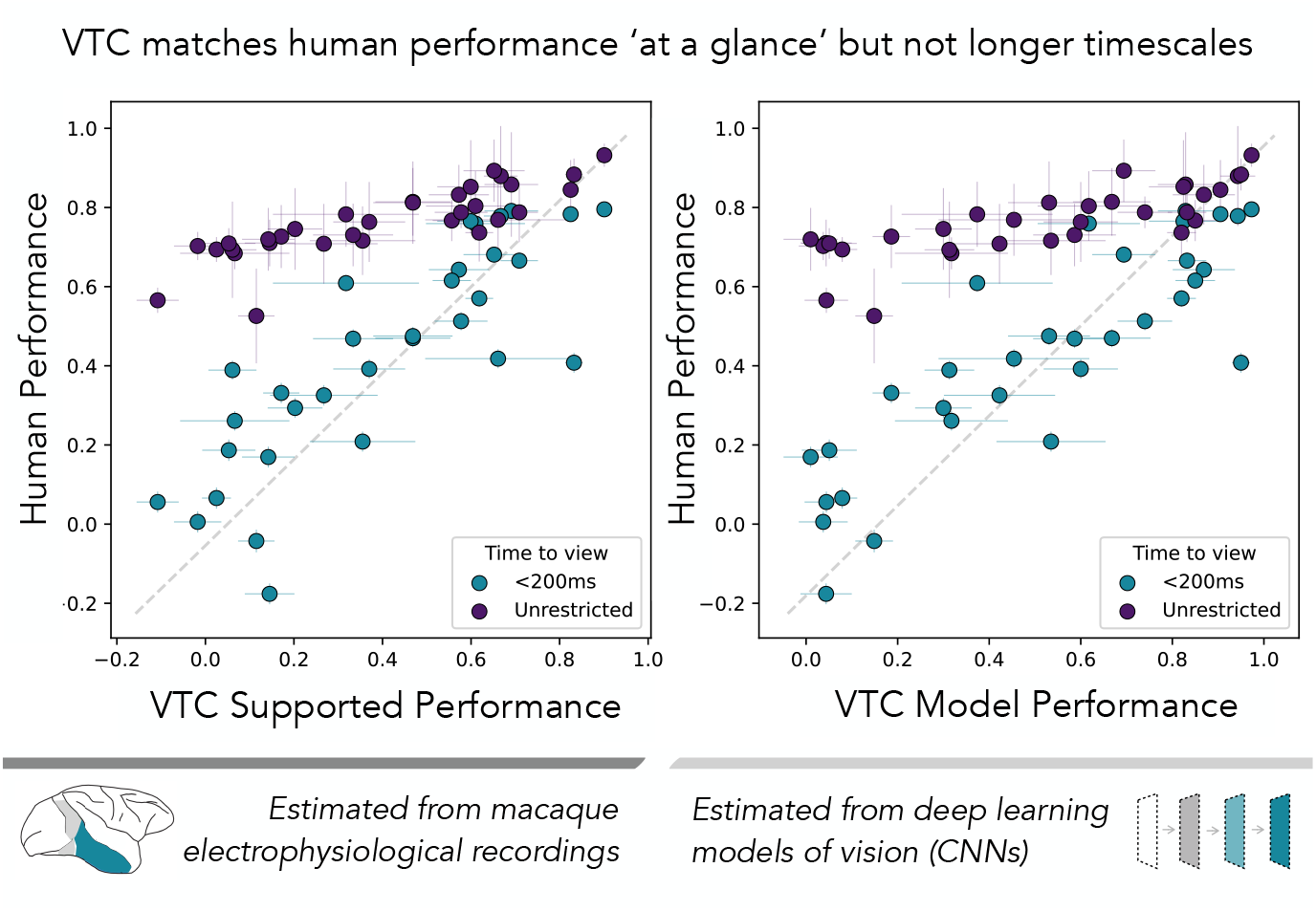
Comparing time restricted/unrestricted human performance to VTC responses. We estimate ‘VTC supported performance’ (left; x axis) from electrophysiological responses collected from high-level visual cortex in the macaque (Majaj et al., 2015). We determine human performance (left; y axis) by administering the same stimuli to participants online. Each point corresponds to the averaged performance across all participants when presented with a single object (i.e., averaging over all trials with the same sample object). The error bars on each point correspond to the standard error of the mean (SEM) over all trials/participants for each object. Using a zero-delay match-to-sample paradigm, we find that when participants are only able to encode the ‘sample’ image for 100 ms, human visual inferences are predicted by VTC supported performance (left; green). When participants can view the same stimuli with unrestricted viewing times, using a simultaneous visual discrimination (i.e., ‘oddity’) task, human visual inferences radically exceed VTC supported performance (left; purple). We demonstrate the same pattern in relationship to a computational proxy for VTC (right): time restricted human visual inferences are predicted by a VTC-like model (right; green), while time unrestricted visual inferences radically exceed VTC model performance (right; purple). Results from all experiments are normalized such that chance performance is plotted at zero.

### 2.4 VTC models exhibit the same pattern of performance as VTC recordings

Next, we estimate ‘VTC Model Performance’ with the same leave-one-out cross-validation strategy used above: we pass the same images which were presented to non-human primates in the Majaj et al., 2015 to a task-optimized convolutional neural network, extract model representations to each image, and then use the same protocol developed to estimate VTC supported performance (Methods: Determining VTC model performance on Majaj et al., 2015). This approach results in a single estimate of VTC Model Performance on each item (*N* = 32). This computational proxy for VTC predicts neural responses in high-level visual cortex (*β* = .81, *F* (1, 30) = 13.33, *P* = 4×10^*−*14^; Fig. 4, and is outperformed by time unrestricted participants (*β* = .16, *t*(31) = 5.38, *P* = 7×10^*−*6^; Fig. 4, right purple). Critically, just as with the relation to VTC supported performance, the relationship between human performance and VTC model performance is evident in the temporal interaction term (*β* = 0.51, *F* (3, 60) = 6.17, *P* = 6 × 10^*−*8^). These data validate the use of these computational models as proxies for VTC and, notably, demonstrate that they display the same pattern of accuracy in relation to time restricted/unrestricted human visual inferences.

### 2.5 Experimental observations are driven by temporal dynamics not design changes

Finally, we demonstrate the difference between time restricted and time unrestricted performance is not a consequence of design changes (i.e., oddity vs. match-to-sample), but the consequence of time restriction. That is, it is possible that by keeping all stimulus present for the duration of each trial, the oddity design might simply alleviate working memory demands that are present in the match-to-sample task. To evaluate this possibility, we administer a ‘self-paced’ match-to-sample task where participants are given unrestricted time to encode the stimulus, before the choose to move on to the match screen. This task retains all of the cognitive demands of the 100ms match-to-sample design, while permitted participants more time to encode the ‘sample’ image. Just as in the oddity task, we find that participants (*N* = 60) in the time unrestricted match-to-sample experiment substantially exceed the performance of time restricted participants (paired ttest *t*(30) = 12.01, *P* = 5.44 × 10^*−*13^; Fig. S9): increasing the time participants can encode the stimulus on the sample screen leads to substantial performance enhancement over time restricted visual inferences. We note that participants in the simultaneous visual discrimination task do outperform participants in the self-paced match-to-sample task (*t*(30) = 3.71, *P* = 0.0008), suggesting that there are other cognitive factors in the oddity design that may contribute to supra-VTC performance (e.g., less working memory demands). These data make clear that humans are able to significantly outperform VTC given longer encoding times, revealing a striking contrast between the perceptual inferences that are possible at different timescales.

## 3 Isolating PRC’s role in temporally extended visual inferences

### 3.1 Leveraging a stimulus set that isolates PRC involvement in visual perception

To investigate the neural substrates that enable these temporally extended visual inferences, we leverage a dataset that contains the performance of PRC-lesioned and -intact human participants (Barense et al., 2007). These stimuli have been shown to reveal profound PRC-related deficits on visual inferences. As such, this dataset enables us to evaluate the effects time restriction alongside the effects of PRC lesions. Before describing the experimental data, we briefly outline the rationale behind these experimental stimuli. First, this stimulus set (example stimuli can be seen in Fig. 3) contains both ‘familiar’ objects that belong to semantically familiar object categories (e.g. common animate and inanimate objects such as cars and animals) and ‘novel’ objects that are not semantically familiar to experimental participants (e.g., ‘greebles’). These ‘familiar’ and ‘novel’ objects are further grouped into two experimental conditions: ‘low-ambiguity’ stimuli serve as control trials, intended to evaluate the relative integrity of VTC structures in these PRC-lesioned participants, as well as the cognitive control systems that are needed to perform the task, while ‘high-ambiguity’ trials are designed to evaluate PRC involvement in visual perception. PRC-lesioned participants have been shown to be selectively impaired on the ‘high ambiguity’ condition while exhibiting no impairment on ‘low ambiguity’ trials, with more pronounced deficits for ‘novel’ stimuli (Barense et al., 2007). Originally, these stimuli were administered via a concurrent visual discrimination (i.e., ‘oddity’) task, with each trail containing 4 images (i.e., 3 ‘typicals’ and 1 ‘oddity’) presented in a 2×2 choice arrays. Here, we administered all four conditions of these stimuli to human participants online (N=105) and in lab (n=58) using the same experimental designs used in the previous time restricted/unrestricted protocols. While our experiments used the same stimuli, we note that the number of stimuli presented in each trial has been modified to be standardized across the oddity and match-to-sample experiments: we use minimal number of images necessary for each trial, which is 3 (i.e., 2 ‘typicals’ and 1 ‘oddity’).

### 3.2 Determining VTC model performance on this PRC-dependent stimulus set

To estimate the performance that would be supported by VTC on stimuli in Barense et al., 2007, we develop a simplified vision of the analysis used in Majaj et al., 2015. While we could train a classifier using left-out images for trials in Majaj et al., 2015, this approach is not suitable for stimuli in Barense et al., 2007: each trial is unique, containing images of objects that are not present in other trials. As such, we use a simplified distance metric to determine model performance. In each trial, we pass each image independently to the model, then select the ‘oddity’ to be the item that has the lowest average correlation to other images. Concretely, we pass the 4 images associated with each trial to the model, then extract model responses from an ‘IT like’ model layer. These layer responses were flattened into length F vectors, resulting in an Fx4 response matrix for each trail. To identify the item-by-item similarity between objects in this trial, were used Pearson’s correlation between items in this Fx4 response matrix, generating an 4×4 (item-by-item) correlation matrix. The item with the lowest mean off-diagonal correlation was the model-selected oddity (i.e., the item least like the others) which we then labeled as either correct or incorrect, depending on its correspondence with ground truth. After repeating this protocol for each trial in the experiment, we computed the average accuracy across all trials. This single value, ‘model performance’, represents the performance that would be expected from a uniform readout of IT. We normalize model performance to fall within the range of 0 (chance) to 1 (ceiling) so that we can compare results from these original oddity decisions (i.e., 4-way, with chance at 25%) with current experiments (which were either 3-way oddity decisions, or 2-way match-to-sample decision, with chance at either 33% or 50%, respectively.

### 3.3 Evaluating the temporal dynamics of PRC-dependent visual inferences

We find that time restricted human performance is predicted by VTC modeled performance (subject-by-condition level correlation with VTC model performance: r = 0.77, *β* = 0.57, *F* (1, 195) = 16.92, *P* = 4 × 10^*−*40^; Fig. 5 left, green) while time unrestricted performance diverges from and exceeds time restricted performance (unpaired ttest between the average performance of time restricted/unrestricted condition-level accuracy for all participants: *t*(327) = 5.67, *P* = 3.04 × 10^*−*8^; Fig. 5 left, purple). Critically, when predicting human accuracy from model performance, we observe a significant interaction with stimulus presentation time (*β* = *−*0.17, *F* (3, 325) = *−*3.47, *P* = 6 × 10^*−*4^). These data resemble the same pattern of performance previous observed from PRC-lesioned/-intact performance on this same stimulus set. That is, just as prior work (Bonnen et al., 2021) determined that PRC-lesioned performance is approximated by VTC model performance and PRC-intact performance exceeds it (subset of experiments from this analysis visualized in Fig. 5 right), we find that time restricted performance is approximated by VTC model performance and time unrestricted performance exceeds it (Fig. 5 left). Taken together, these data suggest that time is a critical component to PRC-dependent visual processing. We note that the time restricted/unrestricted comparison using stimuli from Barense et al., 2007 was conducted using two distinct experimental designs. As with the comparison between human and VTC supported performance above, we again administer a ‘self-paced’ match-to-sample task where participants are given unrestricted time to encode the stimulus, before the choose to move on to the match screen. We find that participants (n=20) in the time unrestricted match-to-sample experiment substantially exceed the performance of time restricted participants (paired ttest *t*(275) = 4.56, *P* = 7.17 × 10^*−*6^; Fig. S10), and that there is again an interaction between human and VTC model performance, as a function of stimulus presentation time (*β* = *−*0.16, *F* (3, 273) = *−*2.76, *P* = 0.006). These results demonstrate that the temporal pattern of human visual inferences, previously observed on stimuli from Majaj et al., 2015, is also evident in these experiments using stimuli from Barense et al., 2007.

**Figure 5:**
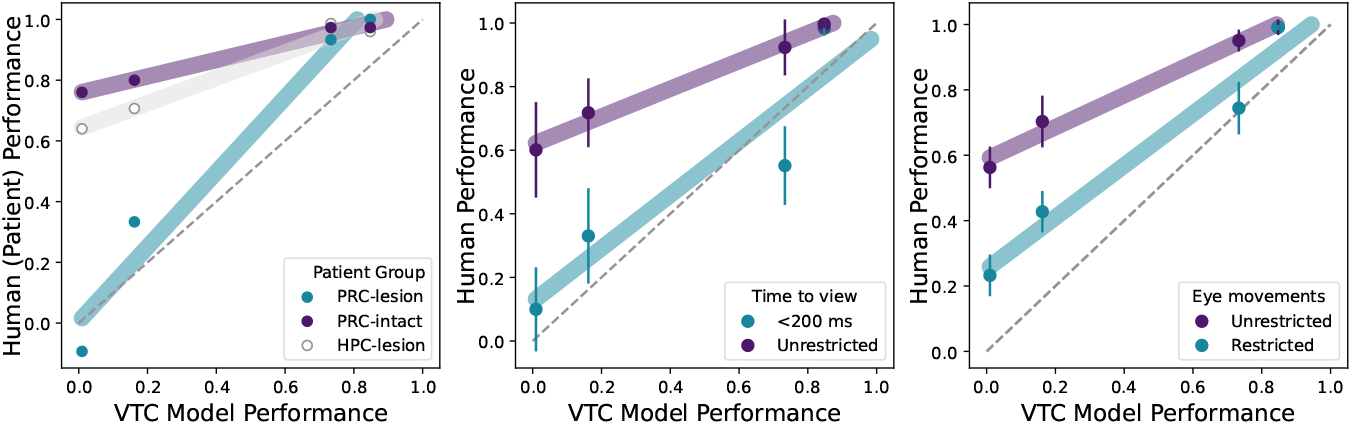
Temporally extended visual inferences rely on PRC and visuospatial sequences. We draw on a set of experimental stimuli from Barense et al., 2007 that reveal the selective contributions of PRC to visual object perception: a computational proxy for VTC predicts the performance of PRC-lesioned participants (left, green) while PRC-intact performance significantly outperform both PRC-lesion participants and VTC models (left, purple). Notably, selective lesions to the hippocampus (left, grey), lead to no such impairment on these visual discrimination tasks. This indicates that these behaviors rely on cortical structures surrounding the HPC, such as PRC. Here, each point corresponds to the averaged performance of all participants, across all trials in one condition. In the current work, we find that the performance of time restricted human participants resembles the accuracy VTC models (middle, green) while time unrestricted performance substantially exceeds time restricted/VTC model performance (middle, purple). Error bars correspond to the standard error of the mean (SEM) over these same trials. When participants are gaze restricted (i.e., required to maintain fixation on a single object location) but given unrestricted viewing time, their performance again resembles VTC modeled accuracy (right, green). Only when participants are allowed to freely move their eyes can they achieve supra-VTC performance (right, purple). Taken together, these data suggest that PRC-supported performance relies on gaze dynamics that are only possible with temporally extended viewing times. Results from all experiments are normalized such that chance performance is plotted at zero.

### 3.4 Time-restriction does not selectively impair PRC-dependent visual inferences

Restricting the time course of visual information processing to *<*200ms appears to affect multiple systems downstream of VTC. For example, PRC-lesioned participants are known to be unimpaired on rotational invariance (Barense et al., 2007). Time restricted performance of neurotypical participants, however, is significantly impaired even on ‘low ambiguity’ (i.e., control) stimuli, when compared to MTL-lesioned patient performance (Fig. 5, left, beneath the diagonal; *t*(13) = *−*6.40, *P* = 2 × 10^*−*5^), presumably because these stimuli are rotated. Indeed, when these stimuli are presented in their canonical orientations in a novel online experiment (*N* = 19), there is no longer a significant difference between time restricted/-unrestricted neurotypical participants (*t*(18) = *−*2.05, *P* = 0.055). That is, restricting viewing times on the sample screen disrupts abilities that appear to be intact in a PRC-lesioned state, as well as those that appear to depend on PRC. As such, time restricted visual inferences do not reflect *selective* disruption of PRC-dependent processes.

## 4 Eyetracking studies to better characterize temporal dynamics

### 4.1 Eye movements are necessary for PRC-dependent visual processing

To evaluate PRC-dependent perceptual dynamics, we conduct a series of in-lab eye tracking experiments (Methods: Eye tracking data collection). Participants’ (*N* = 97) gaze was continuously monitored while they completed an hour-long zero-delay match-to-sample experiment. We first sought to determine whether supra-VTC performance is dependent on longer viewing times per se, or the ability to sequentially sample the visual stimulus via multiple saccades. To evaluate this question, we allow participants to view each stimulus on the sample screen for an unrestricted amount of time (i.e., ‘self-paced’), but restrict their gaze movements: participants must maintain fixation on a single location at the center of each sample image. On the match screen, participants are free to move their gaze, as in previous experiments. Gaze restricted human participants, even with unrestricted viewing time, are significantly impaired relative to participants who could freely view the stimulus on the sample screen (*t*(106) = *−*4.13, *P* = 7.14 × 10^*−*5^). Moreover, the performance of gaze restricted human participants is better approximated by VTC-like models than gaze unrestricted abilities (Fig. 5 center), as is evident in the canonical interaction between human performance and VTC model performance, as a function of gaze restriction (*F* (3, 104) = 5.15, *P* = 1 × 10^*−*6^). Thus, when participants are gaze restricted—because they do not have sufficient time to move their eyes, or because their gaze is fixed—their accuracies resemble the performance supported by a linear readout of VTC. We note that these temporally and foveally restricted accuracies resemble the pattern of impairments following lesions to PRC in human participants (Fig. 5 right, green). Inversely, these data suggest that PRC-supported performance relies on gaze dynamics that are only possible with temporally extended viewing times.

### 4.2 The number of fixations taken by participants predicts PRC dependence

Given the critical role of foveal dynamics in PRC-dependent visual inferences, we next characterize the gaze behaviors at encoding (i.e., viewing the stimulus on the sample screen). First, we evaluate the relationship between encoding time and the number of sequential samples taken by experimental participants, which is given by the number of unique ‘fixations’ on each trial (Methods: Eye tracking data collection). As expected, the duration of time that participants choose to encode the stimulus on the sample screen relates directly to the number of fixations taken on each trial (across all trials *r* = 0.93, *beta* = 2.65, *F* (1, 5803) = 195.35, *P* = 0.000) with a median time spent on each sample screen of 2.77 s, a median number of fixations on each trial of 8, and a median fixation length on each trial of .358 s. We note that, at encoding, there is not a significant difference in the time (across all trials *t*(5803) = 0.90, *P* = 0.36) or number of fixations (*t*(5803) = .80, *P* = 0.42) that participants allocate for trials that rely on PRC from those trials that do not (Fig. 7 left, overlapping box plots). That is, this sequential match-to-sample experimental design enables us to isolate stimulus encoding behaviors from subsequent visual/cognitive operations, such as stimulus-stimulus comparison—as intended. Moving on to the match screen, there continues to be a direct relationship between the time participants view an image and the number of fixations on each trial (*r* = 0.94 *β* = 2.93, *F* (1, 6179) = 224.92, *P* = 0.000), with a median time spent on each sample screen of 2.13 s, a median number of fixations on each trial of 5, and a median fixation length on each trials of 0.45 s. On the match screen, unlike the gaze behavior at encoding, there is a significant difference in the time (*t*(6179) = 38.28, *P* = 5.75 × 10^*−*288^) or number of fixations (*t*(6179) = 36.84, *P* = 9.97 × 10^*−*269^) that participants allocate for trials that rely on PRC from those trials that do not. That is, participants spend more time on PRC-dependent trials (i.e., those conditions were PRC-lesioned participants are impaired, relative to controls; ‘high-ambiguity’ semantically familiar and novel objects; evident in the non overlapping box plots for ‘high’ and ‘low’ ambiguity conditions for both semantically novel and familiar stimuli Fig. 7 right). More explicitly, we note that trials that exhibit greater PRC-lesioned deficits require more time on the match screen (e.g., ‘high ambiguity familiar objects’ vs ‘low ambiguity familiar objects’; *t*(208) = *−* 20.70, *P* = 2.10 × 10^*−*52^; Fig. 7 left, purple) and more sequential visual sampling (e.g., ‘high ambiguity familiar objects’ vs ‘low ambiguity familiar objects’; *t*(208) = *−*20.06, *P* = 1.62 × 10^*−*50^; Fig. 7 left, orange). That is, PRC-lesioned deficits are greater for those trials where participants required more time and fixations. We note that this relationship is evident also in computational proxies of VTC (Fig. 7 right).

**Figure 6:**
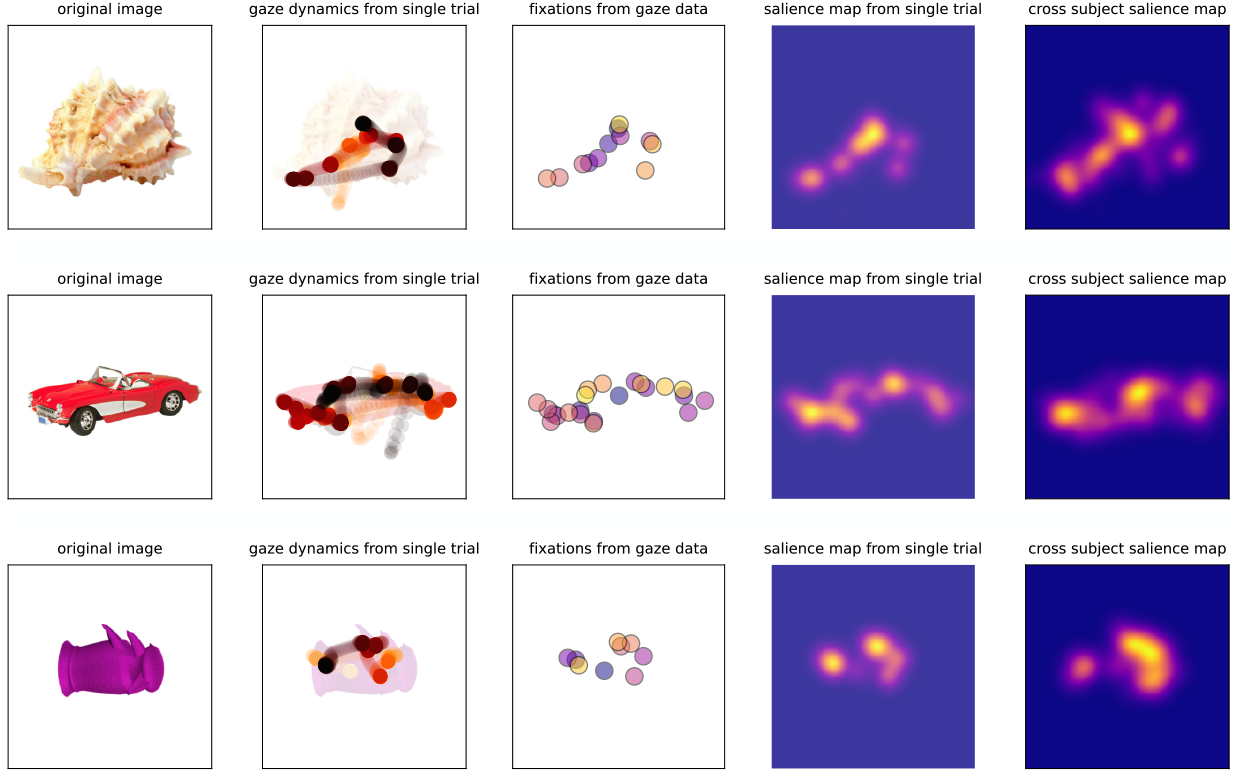
Examples of trial-level gaze data within and across participants. Here we visualize gaze responses to several example images (far left) presented on the sample screen. From the raw time course of gaze data we extract fixations, then generate a subject-level saliency map. For each image, we can also estimate the group-level saliency map by averaging over subject-level maps, where the average number of participants used to estimate group-level saliency map is 16.67 *±* 3.50 STD).

**Figure 7:**
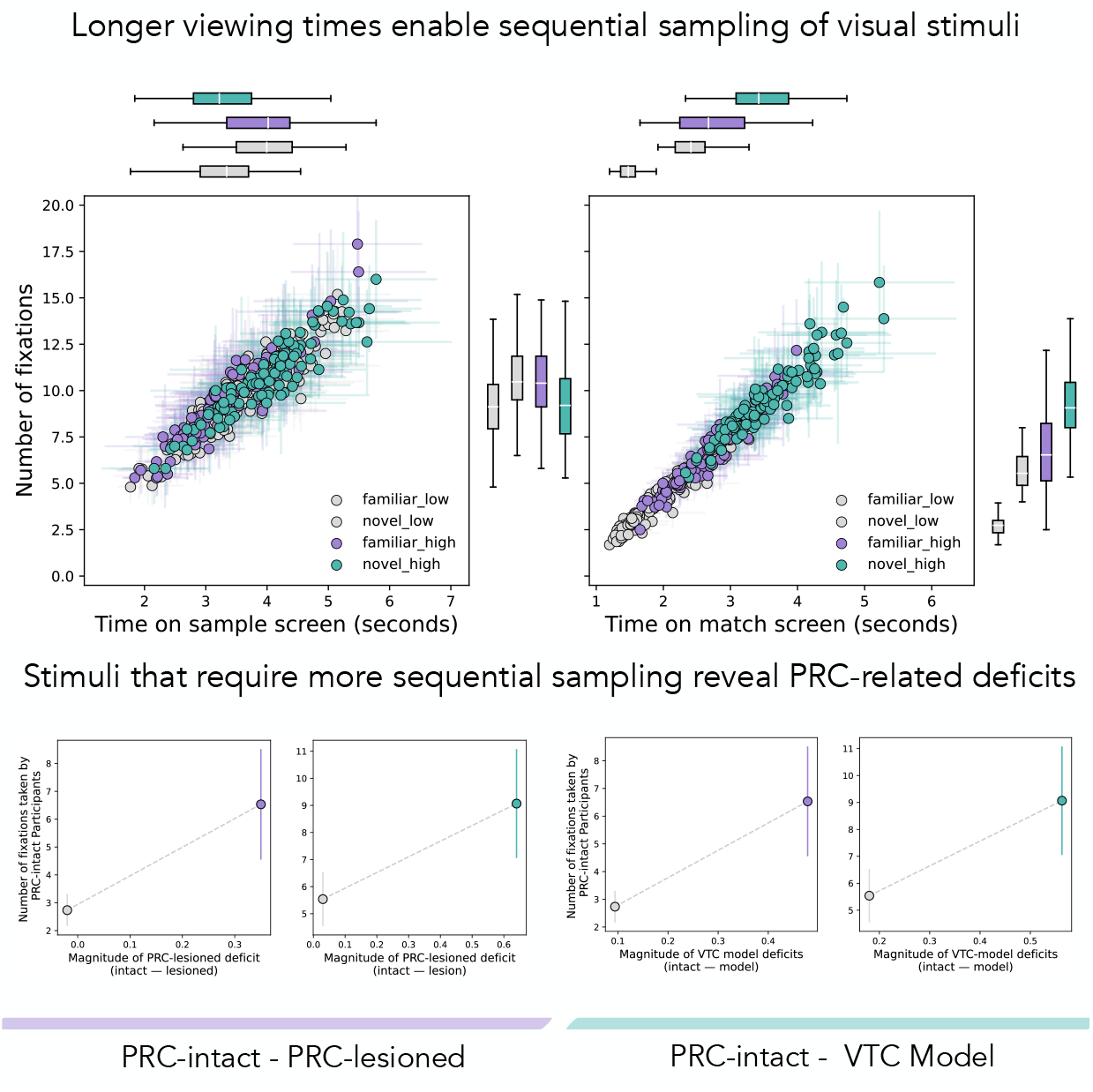
Gaze dynamics of human participants during a PRC-dependent visual task. We administer a sequential match-to-sample task to human participants in lab. We visualize the relationship between reaction time, number of fixations, and stimulus type, for both the sample (top, left) and match (top, right) screens. As intended, on the sample screen, participants are not differentiating between stimulus categories; it is not yet clear what the difficulty of the subsequent match screen will be, and so all images are encoded for roughly the same amount of time (evident in the overlapping box plots between ‘high’ and ‘low’ ambiguity conditions). On the match screen, by comparison, there is a clear distinction between reaction time and number of fixations as a function of stimulus category with the ‘high-ambiguity’ (i.e., PRC-dependent) stimuli requiring substantially more time (non-overlapping boxplots between ‘high’ and ‘low’ ambiguity conditions). That is, this sequential match-to-sample design enables us to isolate stimulus encoding behaviors from subsequent visual/cognitive operations, such as stimulus-stimulus comparison—as intended. We relate these gaze behaviors more explicitly to PRC dependence (bottom). On the x-axis, we plot the magnitude of deficits on each condition, estimated by subtracting the lesion from the intact accuracy and taking the absolute value (i.e., further to the right indicates greater deficits), on the y-axis we plot the average number of fixations taken on the match screen during each trial. We plot this relationship separately for familiar and unfamiliar objects (bottom left, from left to right). These results indicate that PRC-lesioned deficits are greater for those trials where participants required more time and fixations. This relationship is evident also in computational proxies of VTC (bottom, right), where model deficits are defined as the magnitude of different between model performance and neurologically intact participants.

### 4.3 Gaze behaviors are reliable across participants

Finally, we determine whether these gaze behaviors are reliable across participants. We estimate the split-half reliability of these gaze dynamics using the following protocol (Methods: Estimating gaze reliability). First, for each trial, a participant-level salience map is generated from the raw gaze behaviors, smoothed with a Gaussian kernel (example data from a single trial visualized in Fig. 6). For each image, all participant-level salience maps are collected, then randomly partitioned into two splits. The salience maps in each of these random splits are averaged (i.e., across participants), then we determine the correlation between the two (random split-half) salience maps associated with each image. This results in a single numerical value for the random shuffle of participant maps to this image. For each image we repeat this protocol across 100 iterations, storing the correlation across random splits. This results in a distribution of random split-half correlations for each image. To establish an empirical null we compute the correlation between random splits corresponding to the different images within the same trial. Additionally, we estimate the bottom-up salience of each image using standard image processing pipelines (Itti et al., 1998) and also compute the correlation between this bottom-up salience map and the average participant salience maps. Thus, for each image, have three distributions: the split-half correlations between participants’ salience maps, an empirical null (i.e., the correlation between the split half of this image and the split half of another image), and the correlation between participants’ salience maps and a bottom-up salience map. We repeat this process for all images and find that gaze behaviors on each image are remarkably reliable across participants (Fig. 8; purple, median split-half correlation across images of 0.86, STD of 0.095). These image-level reliability scores (Fig. 8 purple, all panels) are significantly greater that the empirical null (unpaired ttest *t*(1258) = 35.97, *P* = 1.90 × 10^*−*195^; Fig. 8 grey, all panels) as well as the fit to bottom-up salience maps (unpaired ttest *t*(835) = 26.38, *P* = 4.58 × 10^*−*112^; Fig. 8 black, all panels), indicating that these gaze behaviors are reliable and not simply driven by bottom-up salience.

**Figure 8:**
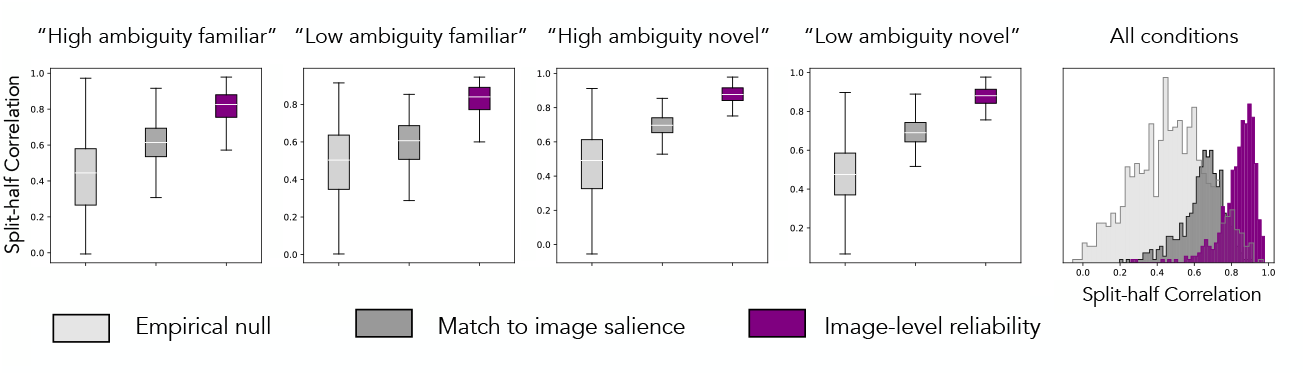
Pattern of spatial attention for each object is reliable across participants. Here we estimate the reliability of item-level gaze behaviors across participants during encoding (i.e., during the sample screen). For each participant, we generate a salience map from the raw gaze behaviors. This yields a salience map for each image a participant observed during the experiment. We can collect all the salience maps for all participants, for a given image, and randomly partitioned these salience maps into two splits. We then averaged across participants in each random split, resulting in two averaged salience maps for this random split. We correlate these two maps, resulting in a single numerical value. We repeat this protocol for each image, storing the correlation values across 100 random splits, resulting in a distribution of random split-half correlations for each image (purple, all plots). We compare this correlation to both an empirically estimated null distribution (light grey, all plots) and the bottom-up salience map of each image (dark grey, all plots). We find that gaze behaviors on each image are reliable across participants. Remarkably, the image-level reliability is greater than the correlation between salience maps and the bottom-up salience estimated from each image. These data indicating that these gaze behaviors are reliable and not simply driven by bottom-up salience.

## 5 Discussion

Our findings indicate that there are complementary neural systems supporting visual object perception. Features that are useful for many downstream behaviors emerge in VTC within 100ms of stimulus presentation (DiCarlo and Cox, 2007), enabling rich visual inferences ‘at a glance’ (Potter and Levy, 1969). However, not all visual inferences are easily (i.e., linearly) separable in this basis space (Jagadeesh and Gardner, 2022). To overcome this limitation, humans sequentially sample visual stimuli via multiple saccades, collecting high-acuity information from multiple task-relevant features. Our work indicates that PRC enables supra-VTC performance by integrating over such visuospatial sequences.

We support this multiple-systems account of perception with computational, electrophysiological, and psychophysics evidence. First, time restricted human performance is predicted by a linear readout of electrophysiological recordings from macaque VTC, as well as a computational proxy for VTC. Conversely, time unrestricted participants substantially outperform a linear readout of VTC. These data indicate that downstream neural structures may be needed to support temporally extended visual inferences. On a second stimulus set we validate these findings: time restricted performance is again predicted by a computational proxy for VTC, while time unrestricted performance exceeds VTC model performance. Through a series of eye tracking experiments we observe that the number of fixations on each image scales linearly with viewing time, and that these patterns of eye movements are both *reliable* across participants and *necessary* for performance. Critically, given prior lesion experiments using this stimulus set, we know that performance depends critically on PRC. That is, PRC enables human participants to outperform a linear readout of VTC on this stimulus set, and these PRC-dependent visual inferences rely on the sequential sampling of images via multiple fixations. This leads us to conclude that PRC enables supra-VTC performance by integrating over visuospatial sequences.

The sequential nature of information processing within medial temporal cortex (MTC) is well recognized in the neuroscientific literature (Buzsáki and Tingley, 2018; Dusek and Eichenbaum, 1997). These sequences are increasingly understood for their ‘compositional’ role in cognitive function (McNamee et al., 2021; Schwartenbeck et al., 2021), enabling animals to flexibly combine learned representations to support novel inferences. The compositional operations of MTC have historically been studied in relation to memory-related functions, where relevant elements are sequentially sampled from prior experience (e.g., Kurth-Nelson et al., 2023). Anatomically, these studies have centered on MTC structures such as hippocampal and entorhinal cortex. Our work extends these findings to the domain of perceptual functions, where the to-be-composed elements are sampled directly from the environment (i.e., the visual stimulus)—or, more concretely, from VTC responses elicited by the environment. These data provide implementation-level support for longstanding theories of visual perception (e.g., Marr, 1982; Ullman, 1987), demonstrating how a finite set of visual features, supported by VTC, can be flexibly repurposed to represent an unbounded set of possible objects, supported by PRC.

These data highlight the computational advantage of integrating neural systems that operate over different timescales. The stimulus properties that VTC can represent are sculpted by the cumulative experience of the organism (Arcaro et al., 2017; Gomez et al., 2019; Nordt et al., 2021; Rajalingham et al., 2020; Srihasam et al., 2012). As such, human performance ‘at a glance’ draws from these amortized visual representations (Dasgupta et al., 2018). This provides a powerful, if limited, basis space of features to represent ongoing experience. In contrast, PRC enables zero-shot performance on ‘out of distribution’ visual inferences—i.e., visual information not present in the information consolidated within VTC, and so not possible ‘at a glance.’ However, consolidating these representations into earlier stages of visual processing provides significant behavioral advantages when these challenges are common occurrences (Goldstone, 1998; Miyashita et al., 1998). That is, visual representations that initially depend on PRC are, over time, consolidated throughout VTC (Erez et al., 2016; Liang et al., 2020; Miyashita et al., 1993). In turn, this enables the elemental features that PRC operates over to be increasingly ‘complex,’ enabling more sophisticated zero-shot visual inferences.

Our work provides algorithmic and architectural constraints for biologically plausible models of human vision. We leave it to future work to develop computational models that leverage these insights to develop more human-like models of vision. Deep learning methods already used throughout the neurosciences (e.g., convolutional neural networks) account for one component of this system; these models have been shown to approximate neural responses and behaviors that depend on VTC, but fail to achieve human-level performance on many visual inferences (Alcorn et al., 2019; Baker et al., 2018; Geirhos et al., 2018; Reizenstein et al., 2021). The divergence between these ‘VTC-like’ models and human performance is often used to critique these computational approaches. Our data support an alternative interpretation: the divergence between humans and ‘VTC-like’ models highlight the multisystem nature of perception. As such, biologically plausible models of human vision should be designed to reflect the algorithmic and architectural constraints we have identified here.

## Acknowledgements

This work is supported by the National Institute of Neurological Disorders and Stroke within the NIH (Award Number F99NS125816) and Stanford’s Center for Mind Brain Behavior and Technology.

## 6 Methods

### 6.1 Estimating VTC supported performance on stimuli from Majaj et al., 2015

In order to estimate the performance supported by VTC and compare to human visual inferences, we utilize stimuli and electrophysiological data from a previous experiment (Majaj et al., 2015). This stimulus set consists of 5760 unique black and white images. Every black and white image contains one of 64 objects, each belonging to one of eight categories, rendered in different orientations and projected onto random backgrounds. This results in 90 images per object. For each image, the original study collected population-level electrophysiological responses recorded from primate V4 and IT. To estimate VTC supported performance on the stimuli above, we use a modified leave-one-out cross validation strategy. For a given sample_*ij*_ trial we construct a random combination of three-way oddity tasks to be used as training data; we sample without replacement from the pool of all images of object_*i*_ and object_*j*_, excluding only those three stimuli that were present in sample_*ij*_. This yields ‘pseudo oddity experiments’ where each trial contains two typical objects and one oddity that have the same identity as the objects in sample_*ij*_ and are randomly configured (different viewpoints, different backgrounds, different orders). These ‘pseudo oddity experiments’ are used as training data. For each of these images, we identify neural responses from high-level visual cortex. We then train an L2 regularized linear classifier to identify the oddity across all (*N* = 52) trials in this permutation of pseudo oddity experiments generated for sample_*ij*_. After learning this weighted, linear readout, we evaluate the classifier on the neural responses to sample_*ij*_. This results in a prediction which is binarized into a single outcome *{* 0 | 1*}*, either correct or incorrect. We repeat this protocol across 100 random sample_*ij*_s, and average across them, resulting in a single estimate of VTC supported performance for each pair_*ij*_.

### 6.2 Generating a novel stimulus set from Majaj et al., 2015

To generate human behavioral experiments we subsample stimuli from Majaj et al., 2015. We reconfigure stimuli from four categories into sets that can be used for both within-category oddity tasks as well as match-to-sample tasks. Each trial is designed to have the minimal configuration of objects (*n* = 3) for these two experimental designs: two of the three objects share an identity (two images of the ‘typical’ object_*i*_, presented from two different viewpoints and projected onto different random backgrounds) and the other is of a different identity (one image of the ‘oddity’, object_*j*_, e.g., two animals, where ‘elephant’ and ‘hedghog’ are object_*i*_ and object_*j*_, respectively). We generate a sample trial_*ij*_ for the pair_*ij*_ of objects *i* and *j* by randomly sampling two different objects from the same category, then sampling two images of object_*i*_ (without replacement) and one image of the oddity of object_*j*_, all with random orientations and backgrounds. We create trials composed of stimuli containing 224 objects. We use this protocol for each of the 224 pair_*ij*_s: we generate 5 random combination of trials from each pair_*ij*_ and fix these trials across all experiments (i.e., trial_*ij*_1, trial_*ij*_2, …, trial_*ij*_5), resulting in (224 × 5) 1120 unique trials. We administer a randomized subset (*N* = 100) of these oddity trials to 297 human participants using a browser-based experimental paradigm implemented in jsPsych (De Leeuw, 2015). We note that, given the truly random experimental generation protocol—and, subsequently, the highly variable nature of the stimuli that comprise each trial—there is no guarantee that a given trial_*ij*_ will contain the information sufficient for a correct response. For example, all the faces in a trial might be rotated out of view, such that the correct oddity can not be determined. To address this, of the 5 stimuli presented, for each of the 224 pair_*ij*_s, we restrict our analysis to 1 trial_*ij*_ which is identified empirically. We select this exemplar for each pair_*ij*_ using a single criterion: the item whose average accuracy (across participants) is closest to the average accuracy measured across all trials (across participants) belonging to other categories. This procedure enables us to exclude outliers (due to, for example, the objects not being fully visible on the viewing screen) while not biasing the results in future analyses. Performance estimates are computed across the population of human participants. In this pooled population of behavior, we compute the reliability by determining the averaged correlation over 1000 randomly shuffled split halves: accuracy was reliable at the category (averaging across all oddity trials in a given category, *r* = .97 *±* .03), object (averaging over all oddity objects for a given typical object, *r* = .69 *±* .07), and image level (average performance on a given trial, averaged across participants, *r* = .24 *±* .05). This effect was even more prominent in the estimates of reaction time at the category (*r* = .99 *±* .01), object (*r* = .91 *±* .02), and image level (*r* = .76 *±* .02).

### 6.3 Determining VTC model performance on Majaj et al., 2015

To estimate VTC model performance we use the same protocol as above, but use a task-optimized convolutional neural network instead of neural recordings. We use a task-optimized convolutional neural network (VGG16, Simonyan and Zisserman, 2014), implemented in tensorflow and pre-trained to perform object classification on a large-scale object classification dataset (Imagenet, Deng et al., 2009). For every image in the Majaj et al., 2015 dataset, we convert this stimulus from greyscale to RGB, then resize it to accommodate model input dimensions (224×224×3). We pass each image to the model and extract responses from a penultimate layer (fc6). Then we repeat the protocol outlined above for estimating VTC supported performance. For each object, we generate a suite of ‘pseudo oddity experiments,’ train L2 regularized linear classifier to predict the oddity in each of these pseudo random trails, and evaluate this linear transform of model responses on the test data.

### 6.4 Determining VTC model performance on Barense et al., 2007

When evaluating performance on all experimental conditions, we situate human behavior in relation to a computational proxy for the primate VTC (Methods: Determining VTC model performance on stimuli from Barense et al., 2007). We employ a method similar to representational similarity analyses common in the neuroscientific literature (Kriegeskorte et al., 2008; Mehrer et al., 2020). For each trial, in each available experiment, the stimulus screen containing N objects is segmented into N object-centered images. Given that each trial in Barense et al., 2007 contains 4 independent images, we pass each image to the model, then extracted model responses from a layer that approximates high-level visual cortex (i.e., a layer that fits electrophysiological responses collected from IT cortex, in the macaque). These layer responses were flattened into vectors, and we use the pairwise Pearson’s correlation between items to generate a 4×4 (item-by-item) correlation matrix. The item with the lowest mean off-diagonal correlation was the model-selected oddity (i.e., the item least like the others) which we then labeled as either correct or incorrect, depending on its correspondence with ground truth. After repeating this protocol for each trial in the experiment, we computed the average accuracy across all trials. This single value represents the performance that would be expected from a uniform readout of high-level visual cortex. Thus, we are able to compare visual abilities across all our experimental designs/datasets within a common metric space: VTC model performance. We note that in each each trial in Barense et al., 2007, we only had access to images in the form presented to experimental participants: a single image containing multiple objects. Thus, for each trial, we segment the stimulus screen containing 4 objects into 4 object-centered images. This required segmenting the image using a K-means clustering approach to automatically identify the centroid of each object, defining a bounding box around each of these centroids, then extracted each object from the coordinates of each bounding box. For both ‘familiar’ experiments, which had images with more irregular dimensions, these stimuli were segmented by splitting the original stimulus screen into quadrants of equal size.

### 6.5 PRC-lesioned participants from Barense et al., 2007

We use stimuli and visualize lesion data (Fig. 5) from four experiments in Barense et al., 2007). Here we include information about these participants for convenience, but refer readers to the original manuscript for more details. In the original manuscript, six amnesic patients with focal brain lesions were tested. Based on structural analyses of critical regions within the temporal lobe using an established rating scale validated against volumetric measures, patients were categorized as follows: (1) cases with selective hippocampal damage (hippocampal group; n = 3) and (2) individuals with broader medial temporal lobe damage, including perirhinal cortex (MTL group; n = 3). Of the three patients in the MTL group (age = 69.77 years; education = 10.33 years), two were viral encephalitis cases and the third had experienced traumatic intracerebral bleeding. Of the three patients in the hippocampal group (age = 45.69 years; education = 14.33 years), one had been diagnosed with viral encephalitis, one had anoxia during status epilepticus, and one experienced carbon monoxide induced hypoxia. For the behavioral experiment, 10 young (age = 48.0 years; education = 14.4 years) and 10 elderly (age = 68.38 years; education = 12.1 years) control subjects were age and education matched to the hippocampal and MTL groups, respectively (all *t <* 1.32, all *p >* 0.21).

### 6.6 Online human data collection

Human experimental data were collected online via Amazon Mechanical Turk and Prolific via experiments were implemented in JsPsych (De Leeuw, 2015). Each experiment began with an instruction phase, which introduced them to the task as well as provided 5 practice trials. This provided an opportunity for participants to acclimate themselves to the task and the controls. Once the experiment began, participants initiated the beginning of each trial with a button press (spacebar), such that they can (effectively) pause the experiment whenever they deem appropriate. This was designed to reduce environmental interference in the experiment. Experiments were designed to be completed in 10 minutes and participants were payed at a rate of roughly $16/hour. In addition, participants were awarded a bonus commensurate with their performance, enabling them to earn up to twice the base pay. In order to ensure that participants were fairly compensated for their time, even in the case of a crowdsourcing platform errors, trial-by-trial data were collected throughout the experiment and stored on a custom server built from a Digital Ocean ‘droplet.’

We administer two related experimental designs. First, we use a 3-way concurrent visual discrimination task commonly used to evaluate the role of PRC in perception (Barense et al., 2007; Buckley et al., 2001; Bussey et al., 2002). This design enables us to determine visual inferences that are possible with unlimited viewing time, as all stimuli remain on the screen for the duration of the trial. On each trial, participants are presented with three images and must identify the image that does not match the other two in terms of object identity (i.e., the ‘oddity’). Participants are given upwards of ten seconds to complete each trial. At any point in this duration participants can select the oddity with a button press (right arrow, left arrow, or down arrow) corresponding to those location on the oddity array. After this button press, participants are given feedback related to their performance on that trial, indicating whether their choice was correct or incorrect. If participants do no press a button in these ten seconds, the trial is marked as incorrect, feedback is given on the screen encouraging them to complete each trial within the allotted time.

To evaluate the effect of time restriction on performance, we use a match-to-sample visual discrimination paradigm common among tasks designed to evaluate VTC supported performance (DiCarlo and Cox, 2007; DiCarlo et al., 2012; Majaj et al., 2015). In each match-to-sample trial, participants first view a ‘sample’ image. After 100 ms of stimulus presentation, this sample image disappears, a noise mask is presented for around 30 ms, then two images are presented on the subsequent screen. Participants are instructed to select the image that ‘matches’ the previous image on the sample screen. Participants again have ten seconds to complete each trial, which is a relatively unrestricted amount of time to complete each trial. If participants do no press a button in these ten seconds, the trial is marked as incorrect, with feedback given on the screen encouraging them to complete each trial within the allotted time. Additionally, we administer ‘self-paced’ match to sample experiments. In these experiments, participants determine how long to view the stimulus on the sample screen (up to 10 seconds). After they press a button to move on, the sample image disappears, a noise mask is briefly presented, and then they make their choice on the match screen in a manner identical to the match-to-sample design introduced above.

### 6.7 Eye tracking data collection

Eye tracking was performed using an infrared video-based eye-tracker at 1000 Hz (Eyelink 1000; SR Research). Stimuli were displayed on a 22.5 inch VIEWPixx LCD display (resolution of 1900×1200, refresh rate of 120 Hz) and responses collected via keyboard. Other sources of light were minimized during data collection. The stimulus on the sample screen was presented at the central field of view and spanned up to 10 degrees of visual angle. This stimulus size was selected such that in order to collect high-acuity visual information from various stimulus locations, participants had to move their eyes (i.e., make a saccade). Stimuli on the match screen were the same size, but presented side by side, offset from the horizontal midpoint of the screen by 10 degrees of visual angle. Each experiment began with gaze calibration, then 5 practice trials to acclimate participants to the experimental setup. Each trial was initiated by the participants and began with participants maintaining fixation at the center of the screen (to perform drift correction at the beginning of each trial). Participants completed each trial at their own pace and there was a brief rest period every 5 minutes. This duration of this rest period was at the discretion of each participant. After this rest period, there was another gaze calibration, after which participants again completed a series of trials at their own pace as described above. For all gaze analyses (e.g., evaluating gaze reliability) we estimate gaze-related events (e.g. fixations) directly from the raw gaze data using a standard python library (REMoDNaV; Nyström and Holmqvist, 2010).

### 6.8 Estimating gaze reliability

We estimate the split-half reliability of in-lab gaze dynamics using the following protocol. First, for each trial, a subject-level salience map is generated from the raw gaze behaviors: a 2D histogram is generated from the raw time series, which is then smoothed with a Gaussian kernel. We note that the results reported in this manuscript are robust to the resolution of the 2D histogram and size of the smoothing kernel. This protocol yields a salience map for each image for each subject. We then generate a random split of subjects and partition the salience maps for a given image using this random subject split. We then average across participants in each random split, which results in two salience maps, each corresponding to the random split of participants allotted to that half. We then estimate the correlation between the two (random split-half) salience maps associated with this image. We repeat this protocol for 100 random split-half permutations (i.e., generating a new shuffle of participants each iteration). For each image, we then have a distribution of split-half correlations which enables us to evaluate how similar participants viewed each image. To establish an empirical null we compute the correlation between random splits corresponding the different images within the same trial. Additionally, we estimate the bottom-up salience of each image (Itti et al., 1998) and compute the correlation between this bottom-up salience map and the random splits associated with each permutation of each image.

## Supplemental Information

**Figure 9:**
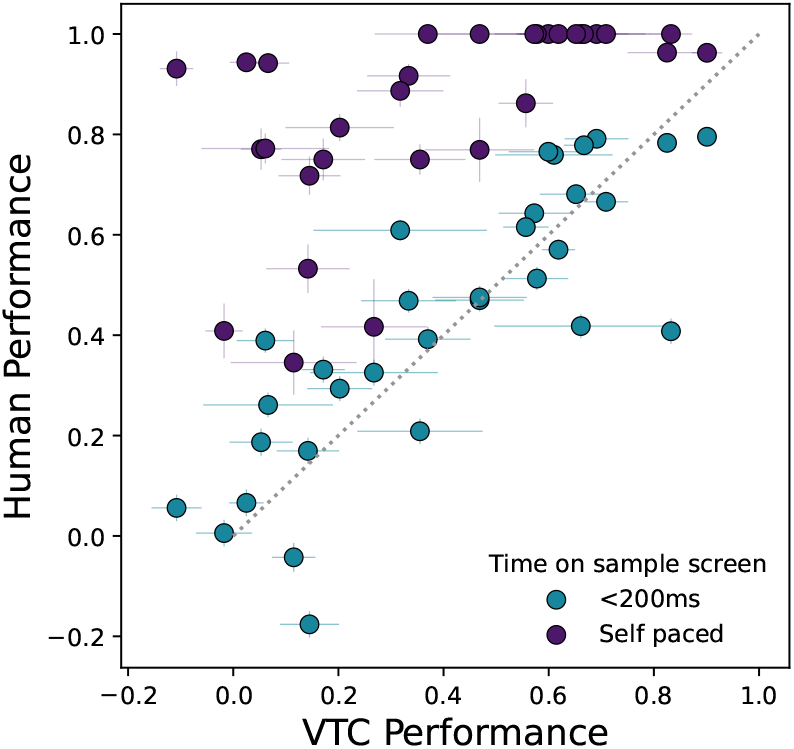
Self-paced match-to-sample performance on stimuli from Majaj et al., 2015

**Figure 10:**
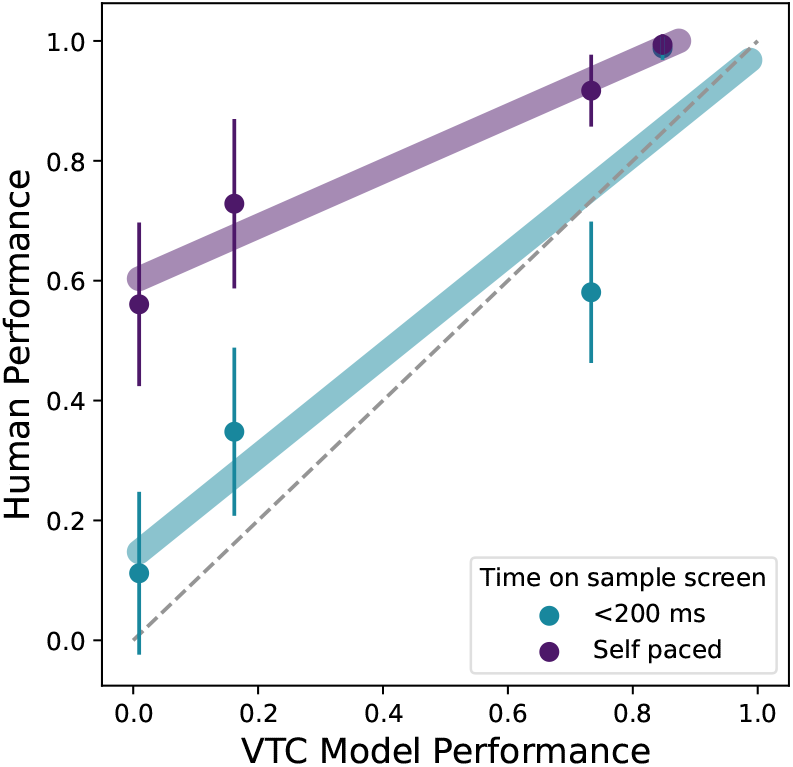
Self-paced match-to-sample performance on stimuli from Barense et al., 2007

